# PAEP and cilia gene splicing changes in endometrial glands during the implantation window in women with recurrent pregnancy loss

**DOI:** 10.1101/2021.09.09.459643

**Authors:** J.E. Pearson-Farr, G Wheway, M.S.A Jongen, P. Goggin, R.M. Lewis, Y. Cheong, J.K. Cleal

## Abstract

Endometrial glands are essential for fertility, consisting of ciliated and secretory cells that facilitate a suitable uterine environment for embryo implantation. This study sought to determine whether an endometrial gland specific transcriptome and splicing profile are altered in women with recurrent pregnancy loss. Our data provide a comprehensive catalogue of cilia and PAEP gene isoforms and relative exon usage in endometrial glands. We report a previously unannotated endometrial gland cilia transcript GALNT11 and its susceptibility to exon skipping. Key endometrial receptivity gene transcripts are also reported to change in endometrial glands of women with recurrent pregnancy loss. The endometrial gland cilia and PAEP targets identified in this study could be used to identify a perturbed endometrium, isolate causes of recurrent pregnancy loss and develop targeted therapies in personalised medicine.

## INTRODUCTION

The endometrium is the lining of the womb in which the embryo implants at the start of pregnancy. It is a heterogeneous tissue containing distinct cell populations encompassing the luminal epithelium, which physically interacts with the embryo, and the glandular epithelium, which produces secretions required for signalling to the embryo and surrounding cells. Hormonally regulated development seeks to ensure that the regenerated endometrium becomes receptive to implantation in the period following ovulation during the window of implantation, the recognised definition of which is days 21-23 of the menstrual cycle (7-10 days after the luteinizing hormone (LH) surge; LH+7-10). Inappropriate receptivity is thought to underlie reproductive failure in women with recurrent pregnancy loss ≥3 miscarriages), subfertility (> 12 months of inability to conceive) and recurrent implantation failure (failure of ≥ 2 IVF cycles with 3 good quality blastocysts replaced). A less receptive endometrium is implicated in subfertility and recurrent implantation failure, as implantation fails in over 50% of IVF patients despite selection of good quality embryos. Whereas an over-receptive endometrium may underlie recurrent pregnancy loss, by allowing the implantation of low quality embryos which increases the risk of pregnancy loss (Macklon & Brosens, 2014). The genetics and cell biology of endometrial receptivity remain poorly understood, limiting scope for clinical intervention in the 2% of women who suffer with unexplained recurrent pregnancy loss (Ford & Schust, 2009).

Studies have attempted to establish molecular markers of endometrial receptivity, yet to date no consistent panel of genes has been identified. Early studies used preselected candidate genes, whereas microarray analysis has identified a potential gene expression profile; termed the endometrial receptivity array (ERA) (Díaz-Gimeno et al., 2011). Analysis using this ERA does suggest a non-receptive endometrium underlies implantation failure (Ruiz-Alonso et al., 2013). A qPCR based panel of selected genes to predict endometrial reactivity has also been developed (Enciso et al., 2018), but again this is not whole genome based. Unfortunately, despite a large amount of data supporting endometrial biomarkers established by omics, there is still little evidence to link omics data to pregnancy outcome (Hernández-Vargas et al., 2020).

Attempts to isolate causes of reproductive failure have been challenged by the cellular heterogeneity of endometrial biopsies (Hu et al., 2014; Suhorutshenko et al., 2018). To date, few studies account for the contribution of different cell types to the overall gene expression profile of the receptive endometrium. To address this, studies need to be carried out on specific populations of cells within the endometrium in order to understand patterns of gene expression in different cells and tissues. Furthermore, to enable early diagnosis in patient care, more work is required to establish a transcriptomic profile of the endometrium from women with a lower order of miscarriages, less distinct than a high order of miscarriages (Craciunas et al., 2021).

Endometrial glands play an essential role in supporting the uterine environment for successful embryo implantation, conceptus development and placentation. Reproductive failure is associated with endometrial gland loss in mouse and sheep gene knock-out models (Filant & Spencer, 2013; Gray et al., 2002), reinforcing the importance of these glandular cell types and their secretions in successful pregnancy. Endometrial gland specific transcriptomic differences are reported in cases of endometriosis (Suda et al., 2018), yet the endometrial gland specific transcriptome has not been investigated in recurrent pregnancy loss. In order to address the gaps in the literature regarding cell-specific expression in different parts of the endometrium, and delineate gene expression in this study, we carried out whole transcriptome RNAseq analysis of endometrial glands from women with recurrent pregnancy loss.

## RESULTS

To investigate differential gene and transcript expression and differential splicing in the endometrial glands of women with recurrent pregnancy loss compared to controls, we performed paired-end 2 × 150 bp RNA sequencing to an average depth of 24.7 million reads per sample on five pairs of isolated endometrial glands from recurrent pregnancy loss patients versus controls matched by the day of the menstrual cycle (Fig 1a). Seventy-three genes were differentially expressed using a 5% FDR in the glandular epithelium from women with recurrent pregnancy loss versus controls (Fig 1b). Of these, 38 genes were upregulated and 35 genes were down regulated in recurrent pregnancy loss. Fifty-seven genes met a more stringent 1.15 log fold change threshold and of these, 24 genes were upregulated and 33 genes were downregulated (Fig 1b). Differential gene expression in the glandular epithelium from women with recurrent pregnancy loss compared to controls included upregulation of the glandular secretory product *PAEP* and *SYT13* involved in transport vesicle docking to the plasma membrane. Significantly enriched biological processes in the glandular epithelium from women with recurrent pregnancy loss included metal ion homeostasis and isoprenoid catabolic processes (Fig 1c). Unsupervised clustering principal component analysis reported that cycle length clustered at day 28 of the menstrual cycle, and therefore was included as a cofounding factor for the differential gene expression analysis.

**Fig 1.**
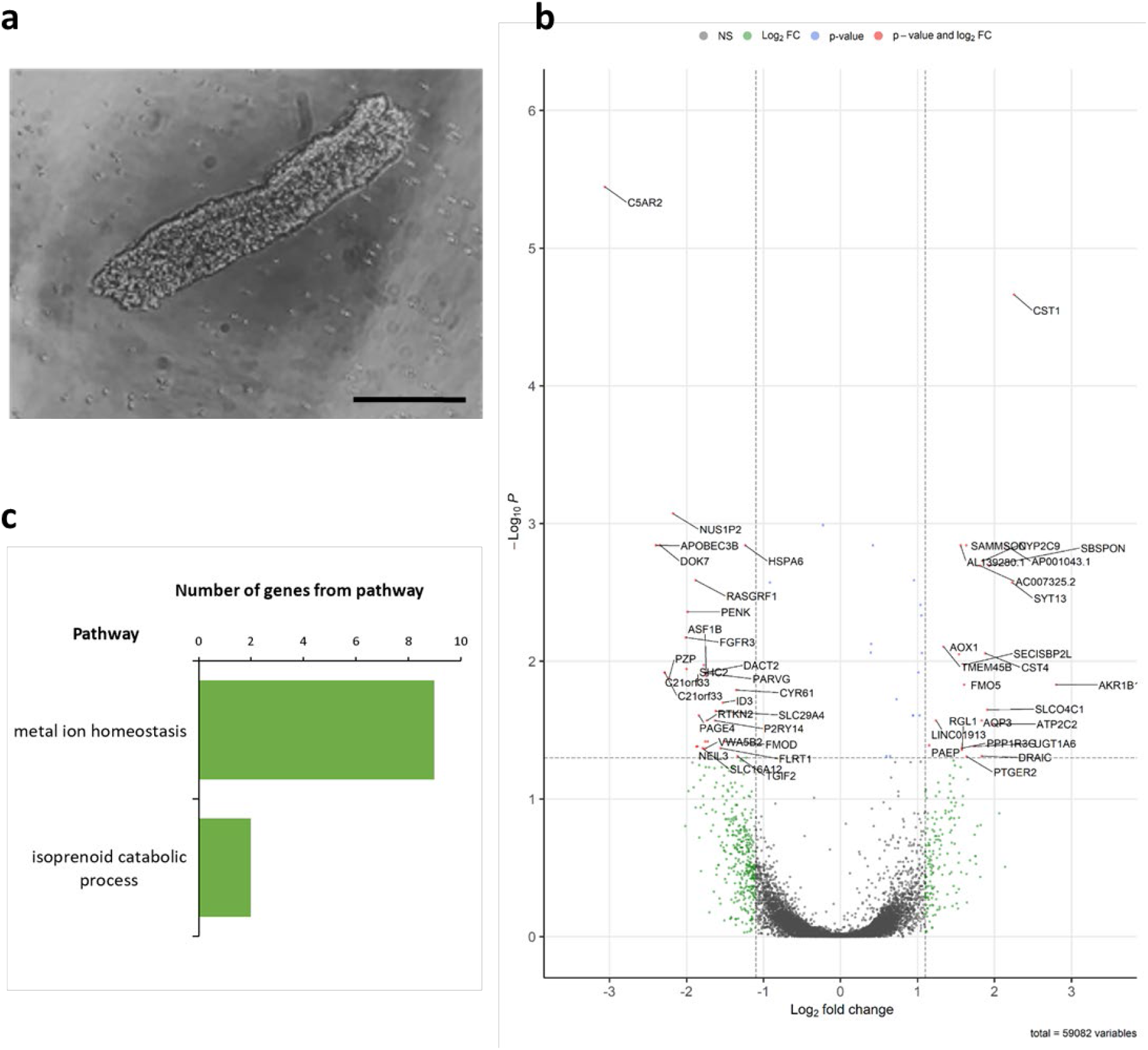
Altered endometrial gland gene expression in women with recurrent pregnancy loss. a) Image of isolated endometrial gland (scale bar = 50 μm) captured using light microscopy, b) Volcano plot representing differentially expressed genes in endometrial glands from women with recurrent pregnancy loss (RPL, n = 5) versus controls (C, n = 5), fold difference between log_2_ normalised expression plotted versus –log_10_ adjusted P-value. c) Biological processes containing differentially expressed genes in endometrial glands from women with recurrent pregnancy loss following analysis of all differentially expressed genes. FDR B&H corrected q value < 0.05.

To further investigate transcriptomic differences between the endometrial glands of women with recurrent pregnancy loss and controls, we carried out transcript-level expression analysis and alternative splicing analysis on our RNAseq data. Two-hundred and seventy-eight differentially expressed gene transcripts were reported in the glandular epithelium of women with recurrent pregnancy loss versus controls. Of those, 257 gene transcripts were upregulated and 21 gene transcripts were downregulated in recurrent pregnancy loss (Fig 2a). Specific gene transcripts included a significant upregulation of pre-identified glandular secretory product *MUC16-204*, glandular progenitor cell marker *LRIG1-205*, intraciliary transport particle *IFT122-204* and known endometrial receptivity marker *LAMB3* (*LAMB-201, LAMB-204* and *LAMB3-203*). The recurrent pregnancy loss group is a heterogeneous group. The two most significantly enriched biological processes in the glandular epithelium of women with recurrent pregnancy loss include tissue morphogenesis and positive regulation of cell differentiation (Fig 2b).

**Fig 2.**
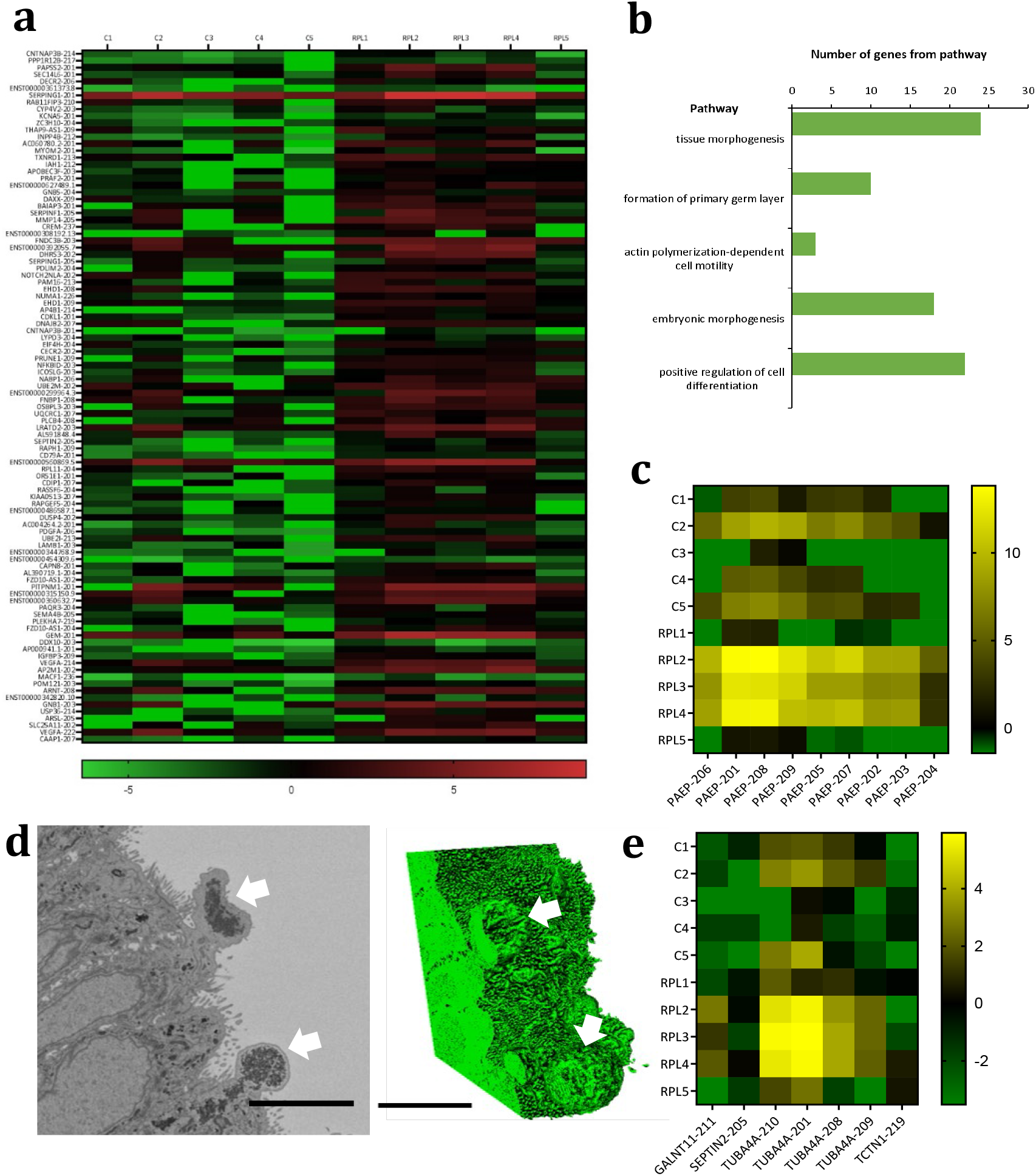
Altered endometrial gland gene transcript expression in women with recurrent pregnancy loss. a) Heat map representing top 100 differentially expressed gene transcripts in endometrial glands from women with recurrent pregnancy loss (RPL, n = 5) versus controls (C, n = 5), data presented as log_2_. b) Biological processes containing differentially expressed gene transcripts in endometrial glands from women with recurrent pregnancy loss following analysis of all differentially expressed genes. FDR B&H corrected q value < 0.05. c) Heat map representing altered PAEP gene transcript expression in endometrial glands in recurrent pregnancy loss versus controls. d) TEM image and 3D reconstruction of electron dense material budding from the apical surface of the glandular epithelium (white arrows), scale bar = 5 μm. e) Heat map representing altered cilia gene transcript expression in endometrial glands in recurrent pregnancy loss versus controls.

Alternative splicing events were significantly different in the glandular epithelium from women with recurrent pregnancy loss versus controls. These included exon skipping, intron retention, mutually exclusive exons, alternative 3’ splice site and alternative 5’ splice site. Four hundred and eighty-five gene transcripts were significantly altered in exon skipping (< 0.05 FDR), and were enriched in the cilium and microtubule skeleton cellular components (Fig 3a). Specific human gene transcripts included those involved in ciliary function (*GALNT11, FBXL13* and *LRRC6*). A previously unannotated *GALNT11* transcript was reported (GALNT11-211), although *GALNT11* is expressed at low levels there was a significant splicing difference between the numbers of reads spanning the exons (Fig 3d). *GALNT11* exon skipped starting at coordinate 152027596 and ending at coordinate 152027718. Other cilia gene transcripts present include *SEPTIN2-205, TUBA4A-210, TUBA4A-201, TUBA4A-208, TUBA4A-209* and *TCTN1-219* (Fig 2e). *GAS5* and *DYNLL1* also had significantly altered intron retention events.

**Fig 3.**
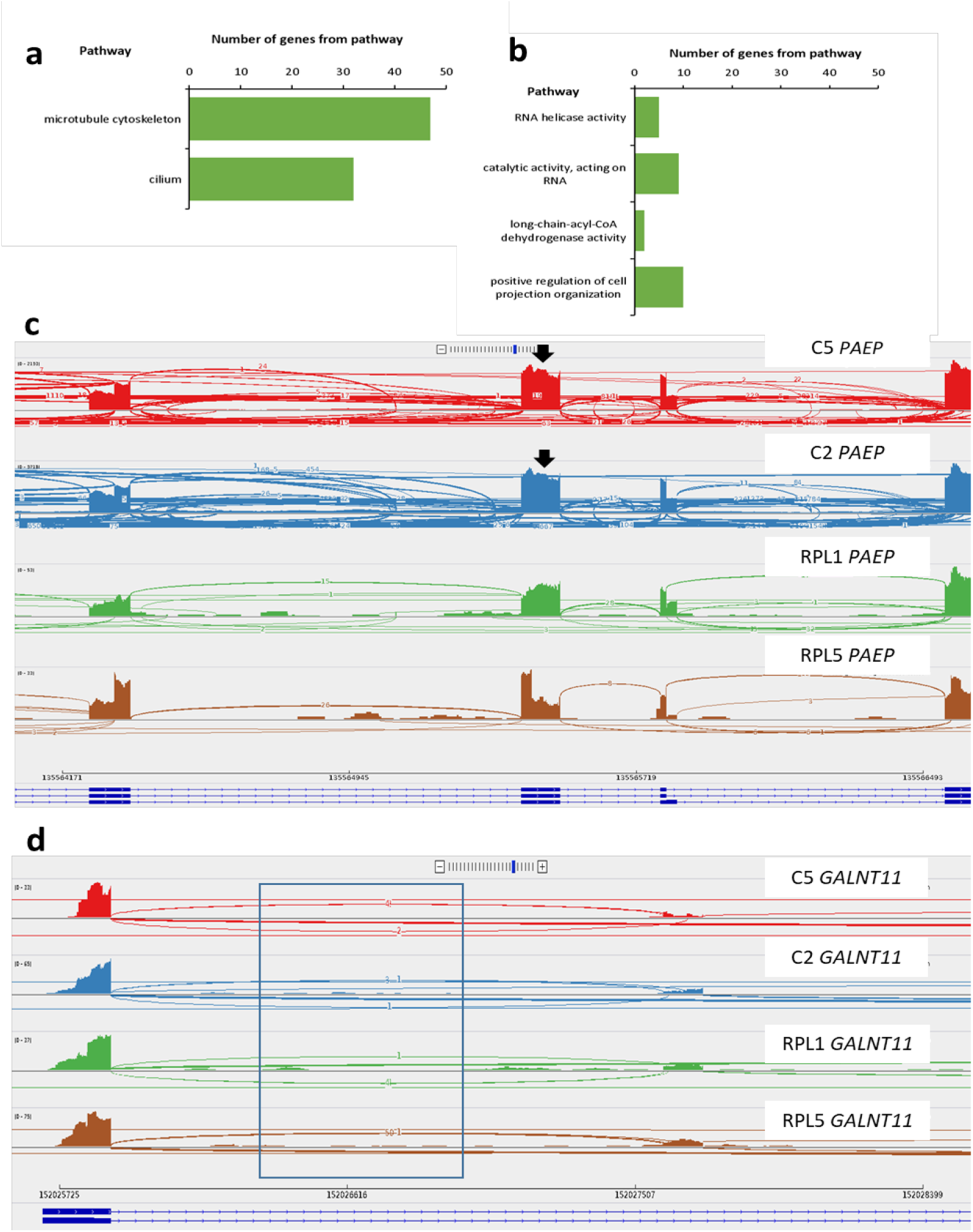
Alternative splicing events in endometrial glands in women with recurrent pregnancy loss. a) Cellular components commonly undergoing exon skipping in the endometrial glands of women with recurrent pregnancy loss (n = 5) compared to controls (n = 5). FDR Bonferroni corrected q value < 0.05. b) Molecular functions and biological process commonly undergoing intron retention in the endometrial glands of women with recurrent pregnancy loss compared to controls. FDR Bonferroni corrected q value. c) Sashimi plots of differentially spliced *PAEP* demonstrating a decline in exon skipping events in recurrent pregnancy loss (RPL; green and brown) compared to controls (C; red and blue; black arrows highlight exon skipping events). Each plot shows gene expression (bar graph), the number of reads split across the splice junction (curved lines), exons (blue bar at the bottom of the plot) and introns of the corresponding gene (dotted lines at the bottom of the plot). d) Sashimi plot demonstrating increased reads split across splice junctions in GALNT11 in women with recurrent pregnancy loss compared to controls.

Eighty-three gene transcripts were significantly changed in intron retention (< 0.05 FDR) and were enriched in biological processes RNA helicase activity, catalytic activity acting on RNA, long chain dehydrogenase activity and positive regulation of projection organization (Fig 3b). Eighty-six gene transcripts had significantly altered mutually exclusive exons in the glandular epithelium of women with recurrent pregnancy loss compared to controls (< 0.05 FDR). Seventy gene transcripts had significantly alternative 3’prime splice sites in the glandular epithelium of women with recurrent pregnancy loss compared to controls (< 0.05 FDR). Finally, 49 gene transcripts had significantly alternative 5’ splice sites in the glandular epithelium from women with recurrent pregnancy loss compared to controls (< 0.05 FDR).

Seven *PAEP* gene transcripts were upregulated in endometrial glands from women with recurrent pregnancy loss compared to controls. These included *PAEP-206, PAEP-201, PAEP-208, PAEP-209, PAEP-205, PAEP-207* and *PAEP-202* (Fig 2c). Alternative splicing of *PAEP* in endometrial glands of women with recurrent pregnancy loss versus controls (<0.05 FDR) demonstrated cryptic splice site usage in *PAEP* (Fig 3c). Three *PAEP* exon skipping events were significantly altered in recurrent pregnancy loss versus controls. Exons skipped included *PAEP* start coordinate 135562359 and end coordinate 135562433, *PAEP* 135562293 -135562433, and *PAEP* 135564243 -135564354. *PAEP* intron retention events were not significantly changed in the glandular epithelium of women with recurrent pregnancy loss versus controls. Five mutually exclusive exon *PAEP* events were significantly different in glandular epithelium from women with recurrent pregnancy loss versus controls (< 0.05 FDR; Table 2). Two *PAEP* alternative 3’ splice sites were significantly altered in the glandular epithelium from women with recurrent pregnancy loss compared to controls. No alternative *PAEP* 5’ splice sites however, were altered in the glandular epithelium from women with recurrent pregnancy loss compared to controls. TEM imaging in endometrial gland secretory cells from control participants, demonstrated structures containing electron dense material budding from the apical surface of the epithelium. SBF SEM reconstructions show that these were approximately spherical in shape (Fig 2d).

**Table 1.** Participant clinical characteristics

| Characteristic | Control (n = 5)<br>mean (SD) | Recurrent<br>pregnancy loss (n =<br>5) mean (SD) |
| --- | --- | --- |
| **Demographic characteristics** |  |  |
| Age | 28.8 (5.1) | 33.0 (3.9) |
| BMI | 22.0 (2.2) | 24.2 (3.0) |
| **Menstrual cycle characteristics** |  |  |
| Day of menstrual cycle | 21.0 (2.0) | 21.4 (1.9) |
| Length of menstrual cycle | 28.8 (0.9) | 27.8 (1.9) |
| **Fertility history** |  |  |
| Contraceptive use in last year | none | none |
| Number of pregnancies | 0.6 (0.5) | 7.2 (1.9) * |
| Number of miscarriages | 0.0 (0.0) | 6.0 (2.5) * |
| Met AMH criteria for egg donation | Yes | n/a |

\* = $p < 0.01$ indicates significantly different from control group, AMH = anti-mullerian hormone, n/a = not applicable.

**Table 2.** Mutually exclusive exons events for PAEP significantly altered in the glandular epithelium of women with recurrent pregnancy loss compared to controls.

|  | Start<br>coordinate<br>1<sup>st</sup> exon | End<br>coordinate<br>1<sup>st</sup> exon | Start<br>coordinate<br>2<sup>nd</sup> exon | End<br>coordinate<br>2<sup>nd</sup> exon |
| --- | --- | --- | --- | --- |
| PAEP | 135565413 | 135565514 | 135565784 | 135565830 |
| PAEP | 135562819 | 135562893 | 135565409 | 135565514 |
| PAEP | 135562819 | 135562893 | 135565337 | 135565514 |
| PAEP | 135562819 | 135562893 | 135565413 | 135565514 |
| PAEP | 135565337 | 135565514 | 135565784 | 135565830 |

## DISCUSSION

Here we show that endometrial glandular ciliated and secretory cells exhibit dysregulation in women with recurrent pregnancy loss versus controls. Differential transcript expression and relative exon usage are reported, identifying novel *PAEP* and *GALNT11* transcripts produced through the use of cryptic splice sites. Novel cilia and secretory RNA targets identified in this study may pose new avenues for therapeutics for better reproductive outcomes.

We demonstrate upregulation of the glandular secretory protein *PAEP* in women with recurrent pregnancy loss, and that this gene is differentially spliced in women with recurrent pregnancy loss. As well as showing overall increased *PAEP* gene expression, women with recurrent pregnancy loss express novel *PAEP* transcripts produced through use of cryptic splice sites. *PAEP* upregulation has been reported by other studies (Burmenskaya et al., 2017), and alternative *PAEP* splicing has previously been found in the female reproductive tract (Garde et al., 1991) but our data provides further detail on the specific *PAEP* transcripts expressed in the endometrial glands, and the presence of novel transcripts produced through cryptic splice site usage in women with recurrent pregnancy loss. This may represent a novel RNA target for therapeutic intervention in recurrent pregnancy loss. Splice-switching oligonucleotides (SSOs) and other antisense oligonucleotides (ASOs) are becoming increasingly used for the treatment of a range of genetic conditions and cancers, and so identification of RNA transcripts which can be targeted with ASOs and SSOs are very promising avenues for treatment (Dhuri et al., 2020).

Post-translational changes to PAEP have been associated with endometrial cancer, supporting evidence that altered PAEP has an impact on endometrial function (Hautala et al., 2020). Since a recent manuscript has shown that PAEP modulates cytotoxic uterine natural killer cells which pose an adverse effect on the growing fetus and subsequent pregnancy loss, it will be important to perform further studies to investigate and characterise the different PAEP isoforms in recurrent pregnancy loss, relevant splicing profiles and if these events are occurring independently (Dixit & Karande, 2020). In the endometrium, PAEP is a major progesterone-regulated glycoprotein. TGFB2 is also associated with hormone response cell secretion regulation and epithelial migration. Endometrial hormonal profiles regulate the menstrual cycle and ovulation, defining the start of the luteal phase. Changes to hormonal responses by the glandular epithelium such as progesterone receptors could be influenced by a decrease in TGIF2 expression, thereby preventing the repression of TGF beta-responsive genes (Snijders et al., 1992). This could be one factor that accounts for cycle length variation seen in the RNA-sequencing data. The genotype TGFB1 has previously been associated with recurrent pregnancy loss (Magdoud et al., 2013).

The enrichment of differentially spliced cilia gene transcripts, including a novel transcript of GALNT11, in women with recurrent pregnancy loss may offer novel insights into the role of endometrial gland cilia in female fertility. The role of cilia in women’s fertility is a poorly understood area, with most work focussing on Fallopian tube cilia function and dysfunction. Studies in this area are confused by the fact that atypical cilia are commonly observed in the human endometrial epithelium as an artefact of the relatively high turnover rate of this tissue caused by menstruation (Denholm & More, 1980). Furthermore, ciliary beat frequency (CBF) in the Fallopian tube, often used as a proxy for ciliary health and function, varies in relation to the stage of menstrual cycle and anatomical location (Lyons et al., 2002), and is affected by environmental factors such as cigarette smoking (Magers et al., 1995), steroids (Mahmood et al., 1998) and reproductive tract infections (Mardh ey al., 1979; Mardh et al., 1976; McGee et al., 1981). Studies of female fertility in inherited genetic conditions associated with cilia (ciliopathies) such as primary ciliary dyskinesia (PCD) have largely been contradictory and inconclusive (Blyth et al., 2008; Goutaki et al., 2016; Raidt et al., 2015). A recent comprehensive study of fertility in males and females with PCD however, found that women with PCD reported infertility at a higher rate than the general population but a miscarriage rate lower than the general population (Vanaken et al., 2017). This study suggested that mutations in specific cilia genes lead to female infertility and our work further suggests that other cilia genes may be differentially spliced in recurrent pregnancy loss, suggesting a complex role for cilia in female fertility.

In conclusion, our data provide a detailed transcriptomic profile of gland specific differences in recurrent pregnancy loss, leading to a more accurate account of where the gene expression changes are occurring. Our transcriptome data provide a comprehensive catalogue of gland-specific target genes, transcripts and splice variants that are altered in recurrent pregnancy loss. Future work should aim to make investigations on isolated endometrial cell populations to account for cell heterogeneity, which will build evidence for therapeutic targets. The exciting development of endometrial organoid culture methods (Brunel et al., 2020) may be a powerful tool to support studies of molecular mechanism alongside study of clinical patient samples.

## METHODS

### Study participants

Participants were recruited for collection of an endometrial biopsy at a tertiary fertility and gynaecology referral centre in Southampton. Participants recruited met study criteria including aged 18-45, no hormonal contraception and no infections. Two participant groups included control participants and recurrent pregnancy loss participants. Control participants (n = 5) were recruited from fertile women who elected to donate eggs having met criteria for egg donation. Recurrent pregnancy loss participants (n = 5) had a history of three or more miscarriages. Control and recurrent pregnancy loss participants were matched by the day of the menstrual cycle to form 5 pairs. Informed written consent was given by all participants, and ethical approval for this study was given by the Isle of Wight, Portsmouth & South East Hampshire Research Ethics Committee (08/H0502/162). Endometrial biopsies were collected using a Pipelle catheter (Stocker et al., 2017) during the window of implantation (LH + 4 -10) and immediately immersed into 50:50 DMEM/ Ham’s F12 nutrient mixture, containing5% streptomycin for endometrial gland isolation.

### Endometrial gland isolation and RNA preparation

Endometrial gland isolation was performed by enzyme digestion within 1 h of tissue collection. Endometrial tissue pieces were minced into smaller pieces before being digested with 0.7 mg/ml type 1A collagenase in 50:50 DMEM/ Ham’s F12 nutrient mixture, containing 5% streptomycin at 37°C for 2 × 15 min intervals with gentle agitation. The digested cell suspension was then passed through a serum gradient to isolate the endometrial gland fraction of the population. The isolated endometrial gland fraction was then passed through a 50 µm sieve to remove other endometrial cell types. Endometrial glands were then stored in 700 µl QIAol lysis reagent at −80°C until RNA extraction. RNA extraction was carried out using the Qiagen miRNeasy extraction kit. The RNA yield was quantified by Thermo Scientific Nanodrop 1000 Spectrophotometer and RNA quality was analysed using an RNA Nano Chip on an Agilent 2100 Bioanalyser (RNA integrity numbers: C1 = 9.4, C2 = 9.2, C3 = 9.5, C4 = 9.4, C5 = 8.9, RPL1 = 7.7, RPL2 = 9.5, RPL3 = 9.0, RPL4 = 9.4, RPL5 = 8.8).

### Library preparation and RNA sequencing

Library preparation was performed using the TruSeq® Stranded mRNA Library Prep kit (Illumina). The final library was quantified by a Roche KAPA library quantification kit (Illumina) and by the Agilent 2100 Bioanalyser. Paired-end RNA sequencing (2 × 150 bp) was carried out on an Illumina NextSeq 550.

### Serial block face scanning electron microscopy

At the time of tissue processing the endometrial pieces were washed twice with 0.1M sodium cacodylate buffer for 10 min. The endometrial tissue pieces were then stained with respective solutions in order: Osmium/ ferrocyanide fixative for 1 h on ice, Thiocarbohydrazide solution for 20 min, 2% Osmium tetroxide for 30 min, 2% Uranyl acetate for 1 h and Walton’s lead aspartate solution at 60°C for 30 min. Between each incubation and after the Walton’s lead aspartate solution all endometrial pieces had x 5 3 min washes in distilled water (Goggin et al., 2020).

The endometrial pieces were dehydrated using 30%, 50% 70% ethanol and 95% ethanol for 10 min respectively, followed by absolute ethanol x 2 for 20 min and acetonitrile for 20 min. Finally, 50:50 acetonitrile: Spurr resin infiltration overnight. The samples were infiltrated with fresh resin for 6 h the following morning, before being polymerised in a final resin change at 60°C for 16+ h. Polymerised blocks were trimmed and thick sections (500µm) cut, stained with toluidine blue and examined with a light microscope (Palaiologou et al., 2020).

Once an endometrial gland was identified, the resin block was trimmed to a frustum with top face approximately 500 µm^2^ including the gland. This sub-block was mounted onto an aluminium pin with conductive glue and splutter coated with gold/palladium. The endometrial gland was imaged by Gatan 3View inside a FEI Quanta 250 FEGSEM microscope at 3.0kV accelerating voltage and a vacuum of 40 Pa. A stack of consecutive images were generated at a constant voxel size of 0.01 × 0.01 0.05 µm. Segmentation and reconstruction were carried out using Amira and Fiji Image J.

### Transmission electron microscopy

Endometrial tissue pieces were collected into 3% glutaraldehyde 0.1 M sodium cacodylate buffer at pH 7.4 and stored at 4°C until tissue processing. At the time of tissue processing, the endometrial pieces underwent washed in 10 min 0.1 M sodium cacodylate buffer at pH 7.4 plus sucrose and 2 mM CaCl_2_ x 2. The endometrial pieces were incubated in 2% osmium tetroxide 0.1 M sodium cacodylate buffer at pH 7.4 for 2 h. Following a brief rinse in distilled water for 30 sec, the endometrial pieces were incubated in 2% uranyl acetate in ddH_2_0 for 30 min, followed by a distilled water rinse for 30 sec.

The endometrial pieces were dehydrated in 70% ethanol and 95% ethanol for 10 min, followed by absolute ethanol 2 × 20 min and acetonitrile for 30 min. Finally, 50:50 acetonitrile: resin incubation overnight. The samples were incubated in fresh resin for 6 h, before being polymerised and encapsulated in fresh resin at 60°C for 16+ h. Thin sections (90 nm) were cut, stained with lead citrate and imaged using Hitachi HT7700 TEM at 100kV.

## QUANTIFICATION AND STATISTICAL ANALYSIS

### Differential gene expression analysis

Raw FASTQ reads were aligned to the human genome via STAR 2.7.3a alignment using human genome 38 and trimmed with trimmomatic. QC was assessed by FastQC v0.11.3. Gene count data were normalised, and differential gene expression analysis was carried out in RStudio R-4.0.3 package DESeq2 v1.30.1 (Love et al., 2014). The Empirical Bayes approach to false discovery rate (FDR) was applied to correct for multiple testing at 5%. Significance was determined by a Wald test and accepted as p ≤ 0.05. To increase stringency, a further log fold change threshold of 1.15 was applied.

### Gene ontology functional enrichment

Genes which were significantly differentially expressed in the endometrial glands of recurrent pregnancy loss patients compared to controls (no log fold change threshold), were mapped to pathways using the publicly available software Toppgene (Division of Bioinformatics, Cincinnati Children’s Hospital Medical Centre). A Bonferroni or B&H FDR were used to correct findings p < 0.05.

### Differential gene transcript expression analysis and splicing analysis

Raw FASTQ reads underwent adapter trimming and quality filtering (reads containing N > 10%, reads where > 50% of read has Qscore<= 5). Paired FASTQ files were aligned to GRCh38 human genome reference using GENCODE v29 gene annotations (Frankish et al., 2019) and STAR v2.6.0a splice aware aligner (Dobin et al., 2013), using ENCODE recommend options (3.2.2 in the STAR manual (https://github.com/alexdobin/STAR/blob/master/doc/STARmanual.pdf). The two-pass alignment method was used, with soft clipping activated.

### Alignment quality control

BAM files sorted by chromosomal coordinates were assessed for saturation of known splice junctions were calculated using RSeqQC v3.0.1 (Wang et al., 2012).

### Alignment to reference transcriptome and transcript level abundance estimates

Salmon tool was used to perform transcript abundance estimates from raw FASTQ files using selective alignment with a decoy-aware transcriptome built from GRCh38 (Patro et al., 2017).

### Differential splicing analysis

rMATs v4.0.2 (rMATS turbo) was used to statistically measure differences in splicing between replicates of wild-type and mutant sequence (Shen et al., 2014). BAM files aligned with STAR v2.6.0a two-pass method with soft clipping suppressed were used as input.

## DECLARATIONS

### Availability of data and material

Data discussed in this study has been deposited on the NCBI&’s Gene Expression Omnibus (Edgar et al., 2002), and are available through the GEO Series accession number GSE GSE183555.

### Code availability

Not applicable

### Author contributions

JP and JS performed sample collection and laboratory analysis. GW, JS and JP performed analysis of transcriptomic data. All authors contributed interpretation all data and the writing of the study. JC, YC and RL initiated, designed and obtained funding for the study.

### Ethics approval

Isle of Wight, Portsmouth & South East Hampshire Research Ethics Committee (08/H0502/162).

### Consent to participate

Informed written consent was given by all participants.

### Consent for publication

Participant information was anonymised.

